# Phylogenetic relationships in the genus *Avena* based on the nuclear *pgk1* gene

**DOI:** 10.1101/351866

**Authors:** Yuanying Peng, Pingping Zhou, Jun Zhao, Junzhuo Li, Shikui Lai, Nicholas A. Tinker, Shu Liao, Honghai Yan

## Abstract

The phylogenetic relationships among 76 *Avena* taxa, representing 14 diploids, eight tetraploids, and four hexaploids were investigated by using the nuclear plastid 3-phosphoglycerate kinase gene(*pgk1*). A significant deletion (131 bp) was detected in all the C genome homoeologues which reconfirmed a major structural divergence between the A and C genomes. Phylogenetic analysis indicated the C_p_ genome is more closely related to the polyploid species than is the C_v_ genome. Two haplotypes of *pgk1* gene were obtained from most of the AB genome tetraploids. Both types of the *barbata* group showed a close relationship with the A_s_ genome diploid species, supporting the hypothesis that both the A and B genomes are derived from an A_s_ genome. Two haplotypes were also detected in *A. agadiriana*, which showed close relationships with the A_s_ genome diploid and the A_c_ genome diploid, respectively, emphasizing the important role of the A_c_ genome in the evolution of *A. agadiriana.* Three homoeologues of thepgK1 gene were detected in five hexaploid accessions. The homoeologues that might represent the D genome were tightly clustered with the tetraploids *A. marrocana* and *A. murphyi*, but did not show a close relationship with any extant diploid species.

## Introduction

The genus *Avena* L. belongs to the tribe Aveneae of the grass family (Poaceae). It contains approximately 30 species [1–4] reflecting a wide range of morphological and ecological diversity over the temperate and subtropical regions [5]. The evolutionary history of *Avena* species has been discussed for decades, and remains a matter of debate despite considerable research effort in this field. Cytologically, three ploidy levels are recognized in the genus *Avena*: diploid, tetraploid, and hexaploid, with a base number of seven chromosomes [6, 7]. The diploids are divided clearly into two distinct lineages with the A and C genomes. All hexaploid species share the same genomic constitution of ACD, corroborated by fertile interspecific crosses among each other, as well as by their similar genome size[8]. With less certainty, the tetraploids have been designated as AB or AA, AC or DC, and CC genomes [9]. It is noteworthy that the B and D genomes within the polyploid species have not been identified in any extant diploid species. There are three C genome diploid species, which have been grouped into two genome types (C_p_ and C_v_) according to their karyotypes [10]. Both types show a high degree of chromosome affinity to the polyploid C genome [9–14], but none have been undisputedly identified as the C genome progenitor of the polyploids.

The A genome origin of polyploid oats has also been under intense scrutiny. However, there is no conclusive evidence regarding which the A genome diploid contributed to the polyploid oats. There are up to 12 species designated as A genome diploids. These species have been further subdivided into five sub-types of A_c_, A_d_, A_l_, A_p_ and A_s_ genomes, according to their karyotypes [6, 7]. Most research based on karyotype comparisons [6, 15], in situ hybridization [11, 16–18], as well as the alignments of nuclear genes [13, 14] suggest that one of the A_s_ genome species may be the A genome donor of polyploid oats. Alternatively, some studies have proposed the A_c_ genome diploid *A. canariensis* [19], or the A_l_ genome diploid *A. longiglumis* [9, 12] as the most likely A genome donor.

The absence of diploids with the B and D genomes complicates the B and D genome donor identification. It is generally accepted that both B and D genomes are derived from A genomes, due to the high homology between the B and A genomes [11, 20], as well as between the D and A genomes [16, 19, 21]. Our recent study based on high-density genotyping-by-sequencing (GBS) markers [9] provided strong evidence that the three tetraploid species formerly designated as AC genomes are much closer to the C and D genomes of the hexaploids than they are to the hexaploid A genome. These findings suggest that the hexaploid D genome exists in the extant tetraploids. However, no extant diploid species, even the A_c_ genome diploid *A. canariensis,* which was considered as the most likely D genome progenitor based on direct evidence from morphological features [22] and indirect evidence from fluorescent in situ hybridization (FISH) [18], showed enough similarity to the D genome of tetraploid and hexaploid oats to warrant consideration as a direct D genome progenitor.

In the case of the B genome, an initial study of chromosome pairing of hybrids between the AB genome tetraploids and the A_s_ genome diploids suggested that the B genome arose from the A_s_ genome through autoploidization [23]. Recently, another GBS study [19] showed that the AB genome tetraploid species fell into a tight cluster with A_s_ genome diploids, also supporting the hypothesis that the B genome arose through minor divergence following autoploidization. However, other evidence from C-banding [24], FISH [17], RAPD markers [25], and DNA sequence alignment [14] has indicated a clear distinction between A and B genomes, suggesting an allotetraploid origin of the AB genome tetraploid species. The most probable A genome progenitor of the AB genome tetraploids is assumed to be an A_s_ genome diploid species, while the B genome of these species remains controversial.

Single or low copy nuclear genes are widely used in phylogenetic analyses due to their bi-parental inheritance and to the informativeness of mutations. Such studies have successfully revealed multiple polyploid origins, and clarified hybridization events in a variety of plant families [26, 27]. In a previous study [14], we investigated the relationships among *Avena* species by sequencing the single-copy nuclear acetyl-coA carboxylase gene (*Acc1*). The results provided some useful clues to the relationships of *Avena* species.

The *pgk1* gene, which encodes the plastid 3-phosphoglyceratekinase, is another nuclear gene that has been widely used to reveal the evolutionary history of the *Triticum/Aegilops* complex due to its single copy status per diploid chromosome in grass [26, 28, 29]. The *pgk1* gene is now considered to be superior to the *Acc1* gene in phylogenetic analysis,since it has more parsimony informative sites than the *Acc1* gene [26, 29]. In the present study, we sequenced cloned *Pgk1gene* copies from 76 accessions representing the majority of *Avena* species, in an attempt to further clarify evolutionary events in this important genus.

## Materials and Methods

### Plant materials

A total of 76 accessions from26 *Avena* species were investigated to represent the geographic range of six sections in *Avena,* together with one accession from *Trisetopsis turgidula* as a functional outgroup (Table 1). All seeds were provided by Plant Gene Resources of Canada (PGRC) or the National Small Grains Collection, Agriculture Research Service, United States Department of Agriculture (USDA, ARS) with the exception of the three accessions of *A. insularis,* which were kindly provided by Dr. Rick Jellen, Brigham Young University, Provo, UT, USA. The species *A. atherantha, A. hybrida, A. matritensis* and *A. trichophylla* described in Baum’s [1] monograph and *A. prostrata* described by Ladizinsky [30] were not included due to a lack of viable material.

**Table 1.**
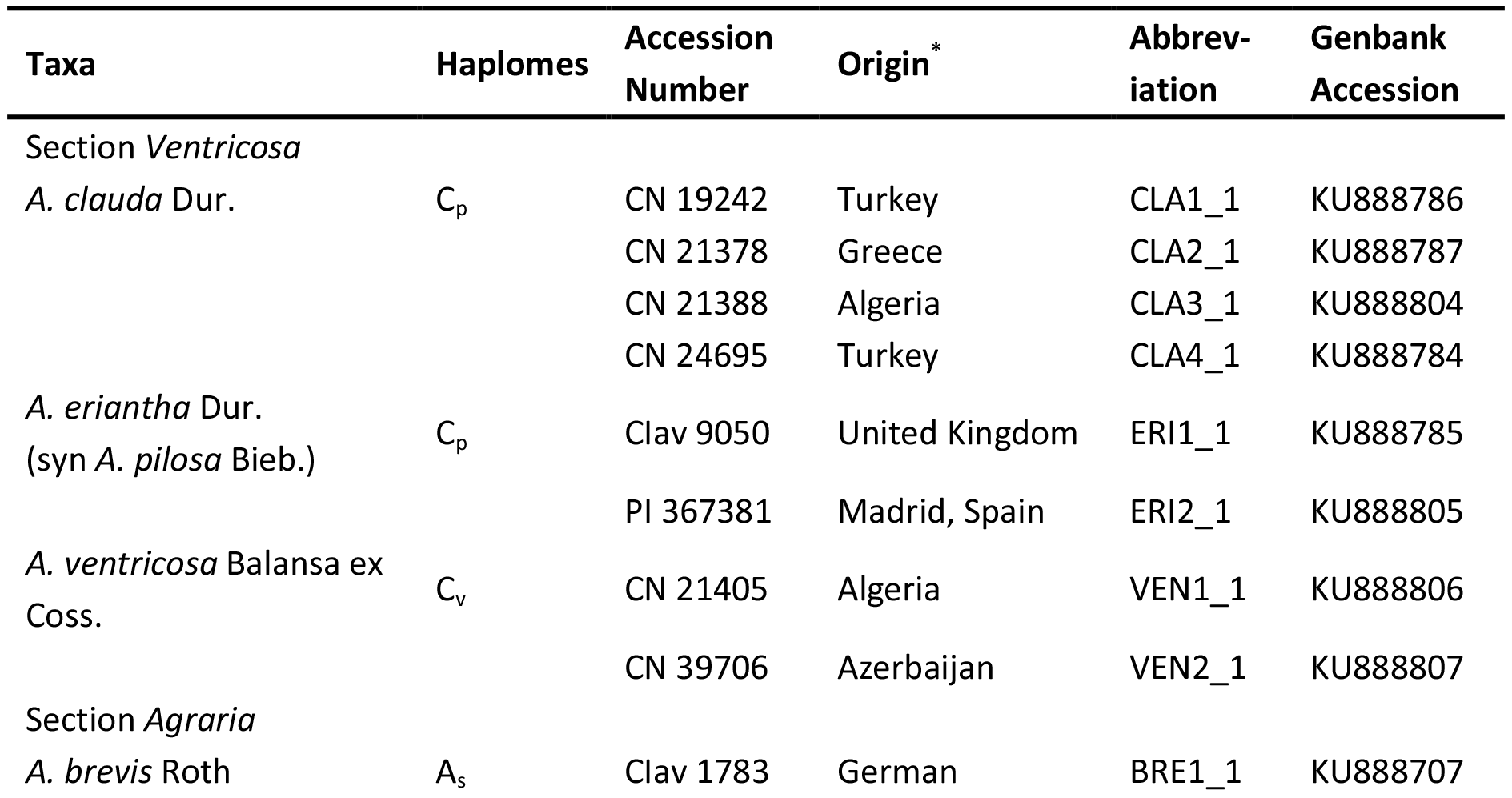
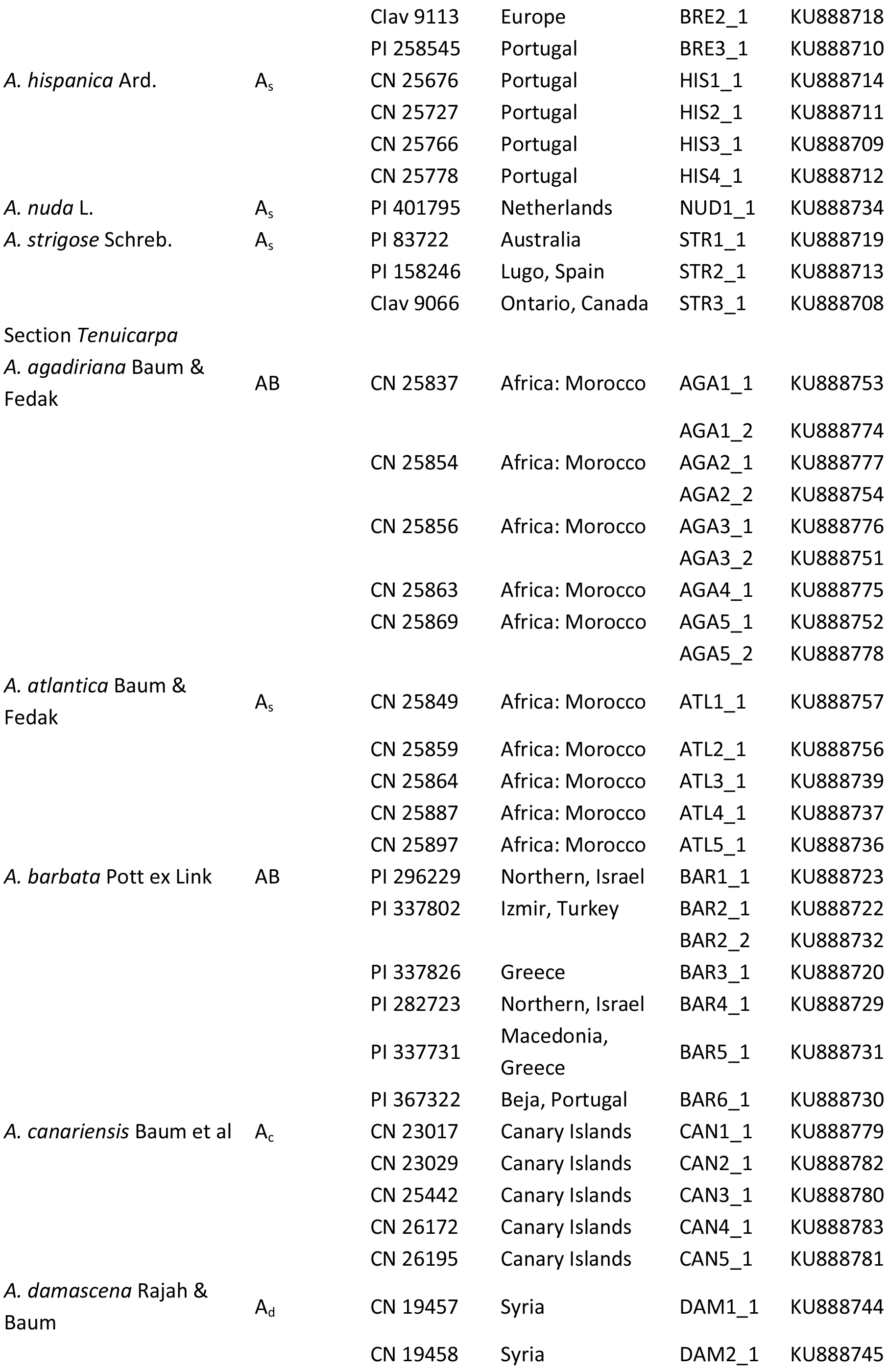
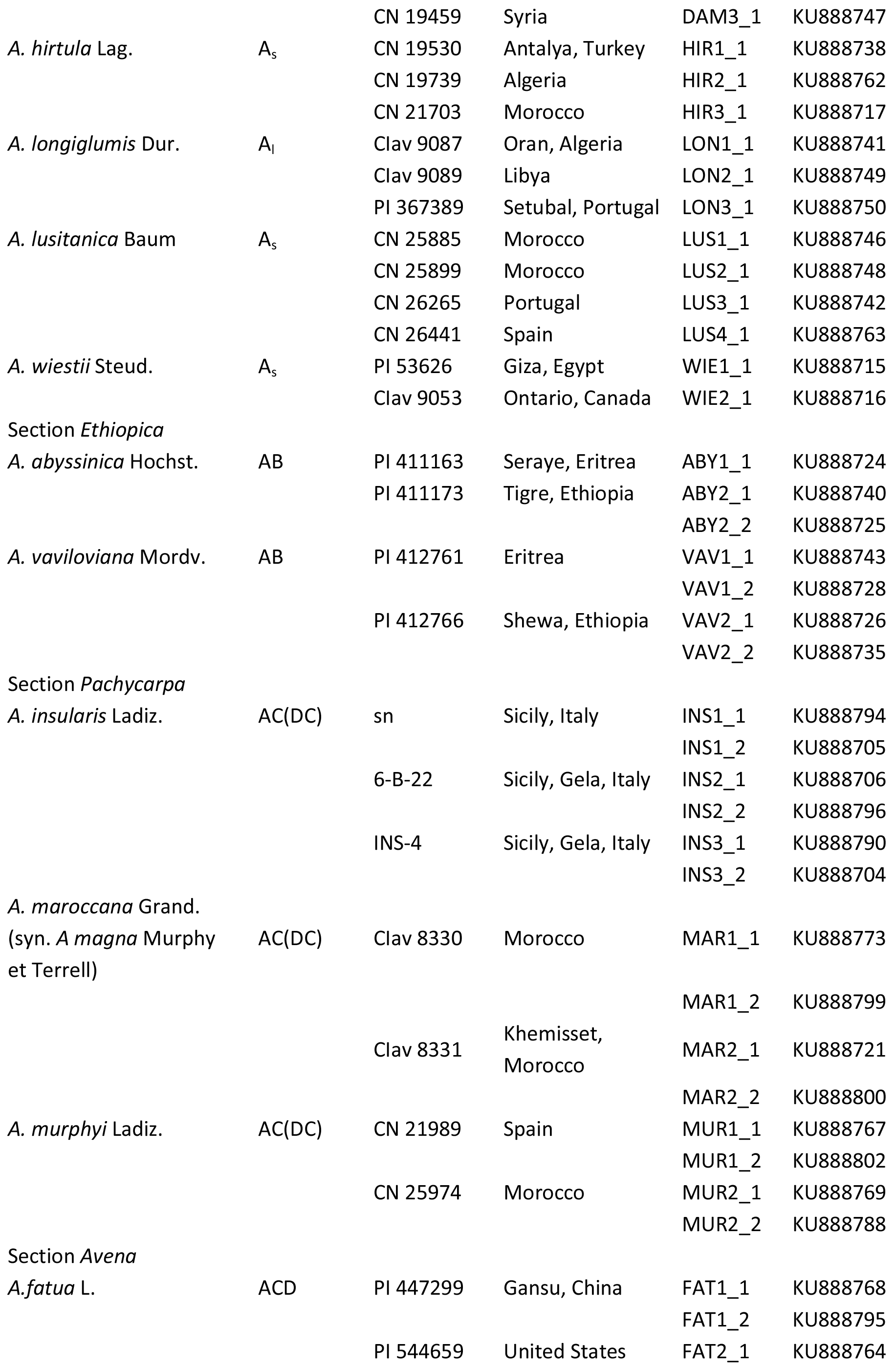
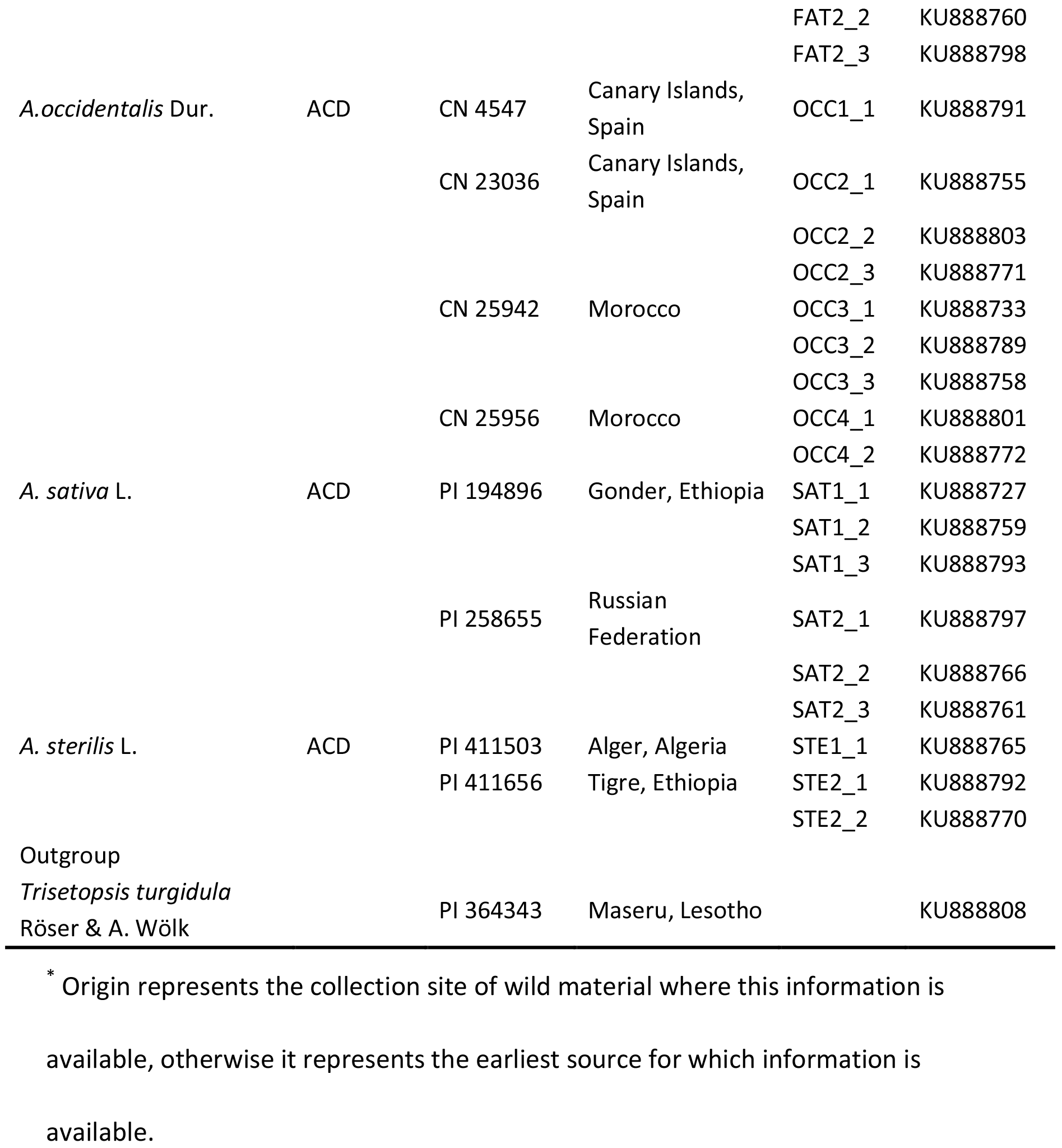
List of materials used in the present study including species, haplomes, accession number, origin, abbreviation displayed in MJ network, and the sequence number in Genbank (https://www.ncbi.nlm.nih.gov).

### DNA isolation, cloning and sequencing

Genomic DNA was isolated from fresh leaves of single plants following a standard CTAB protocol [31]. *pgk1* gene sequences were amplified by using a pair of *pgk1*-specific primers, PGKF1 (5′-TCGTCCTAAGGGTGTTACTCCTAA-3′) and PGKR1 (5′-ACCACCAGTTGAGATGTGGCTCAT-3′) described by Huang et al. [28]. Polymerase chain reactions (PCR) were carried out under cycling conditions reported previously [26]. After estimating the size by 1.0% agarose gel, PCR products were purified using the QIAquick gel extraction kit (QIAGEN Inc., USA). The purified products were cloned into the pMD19-T vector (Takara) following the manufacturer’s instructions. Initially, 6-8 positive clones from each of four accessions from 4 diploid species,including *A. canariensis* (A_c_), *A. longiglumis* (A_l_), *A. strigosa* (A_s_), and *A. clauda* (C_p_), were sequenced to confirm that the *pgk1* gene was present in *Avena* diploid species as a single copy. After confirming its single copy status in diploid species, 2-3 positive clones were selected and sequenced from each accession of the remaining diploid species. In order to isolate all possible homoeologous sequences in polyploid species, 4-6 positive clones from each accession of the tetraploid species and 5-10 positive clones from each accession of the hexaploid species were selected and sequenced. All the cloned PCR products were sequenced on both strands by a commercial company (Sangon Biotech Co., Ltd., Shanghai, China) based on Sanger sequencing technology.

### Sequence alignment and phylogenetic analysis

The homology of sequences was verified usingthe BLAST program in NCBI. In order to reduce the matrix size of the dataset, redundant sequences were removed, keeping one representative sequence if several identical sequences were derived from the same accession. Sequences were aligned using ClustalW software with default parameters [32] followed by manual correction. Substitution saturation of *pgk1* sequenceswasexamined using DAMBE version 5 [33] by calculating and plotting pairwise rates of transitions and transversions against sequence divergence under the TN93 model. Phylogenetic trees were created by using Maximum parsimony (MP), and Bayesian inference (BI). MP analysis was performed on PAUP* 4.0b10 [34] using the heuristic search with 100 random addition sequence replicates and Tree Bisection-Reconnection (TBR) branch swapping algorithms. Bootstrapping with 1000 replicates was estimated to determine the robustness of formed branches [35]. Gaps in the sequence alignment were disregarded using the option ‘gapmode=missing’, which is consistent with an assumption that insertion/deletion events are an independent stochastic process from SNP substitutions. BI analysis was carried out by using MrBayes v3.2 [36]. The best-fit substitution model for BI analysis was GTR+Γ+I, which was determined by using MrModelTest v2.3 under Akaike information criteria (AIC) (http://www.ebc.uu.se/systzoo/staff/nylander.html). Four Markov chain Monte Carlo (MCMC) chains with default priors settings were run simultaneously. To ensure the two runs converged onto the stationary distribution, 6,000,000 generations were run to make the standard deviation of split frequencies fall below 0.01. Samples were taken every 100 generations. The first 25% samples from each run were discarded as the “burn-in”. The 50% majority-rule consensus tree was constructed from the remaining trees. Posterior probability (PP) values were used to evaluate the statistical confidence of each node.

### Network analysis

The median-joining (MJ) network [37] method has been demonstrated to be an effective method for assessing the relationship in closely related lineages [38], and thus was applied in this study. As MJ algorithms are designed for non-recombining molecules [37], DNA recombination was test by using a pragmatic approach-Genetic Algorithm Recombination Detection (GARD), described by Pond et al. [39]. The test was carried out on a web-based interface for GARD at http://www.datamonkey.org/GARD/. Building upon this test, the intron data was used for MJ reconstruction due to the absence of recombination signal, while potential recombination signals were detected in the exon regions. The MJ network analyses was performed using the Network 4.6.1.4 program (Fluxus Technology Ltd, Clare, Suffolk, UK).

## Results

### Sequence analysis

A total of 237 clones were sequenced from 76 accessions of 26 *Avena* species. Following removal of the redundant sequences within each accession, 104 sequences were identified, including one from each of the 44 diploid accessions, 37 unique sequences from 22 tetraploids, and 23 from 10 hexaploids. Theoretically, 44 homoeologues should be isolated from 22 tetraploid accessions, and 30 single-copy homoeologues were expected from 10 hexaploid accessions. However, the full number of expected homoeologues were not isolated from every polyploid species for various potential reasons. In particular, within the AB genome tetraploid species *A. barbata,* only one copy was detected in five of its six accessions, whereas two very similar (only one site varied in exon 2) copies were detected in the sixth accession. This also happened in the hexaploid species *A. sterilis,* for which two accessions provided only two homoeologues each. For these taxa, the missing genome type might be detected by screening a larger number of positive clones, but it is also possible that these accessions contain genomes of high similarity or autopolyploid origin. Another possibility that cannot be ruled out within the polyploids is the loss of one gene copy through homoeologous recombination or deletion.

All of the *pgk1* gene sequences isolated in this study contain 5 exons and 4 introns, covering a total length from 1391 bp to 1527 bp, which is consistent with previous studies of this gene in wheat [28]and *Kengyilia* [26]. The alignment of *pgk1* sequences including both exons and introns resulted in a matrix of 1539 nucleotide positions, of which 11.6% (179/1539) were variable, and 10.1% (155/1539) were parsimony informative. The nucleotide frequencies were 0.264 (A), 0.304 (T), 0.199 (C), and 0.232 (G). A significant (131-bp) insertion/deletion feature (Fig 1A) occurred at position 968, whereby all non-C genome type sequences contained the inserted (or non-deleted) region. Further analysis indicated that this region is likely an inserted inverted repeat, which belongs to the MITE stowaway element. Its secondary structure is shown in Fig 1B. This insertion/deletion event could be used as a genetic marker for rapid diagnosis of *Avena* species containing the C genome.

**Fig 1.**
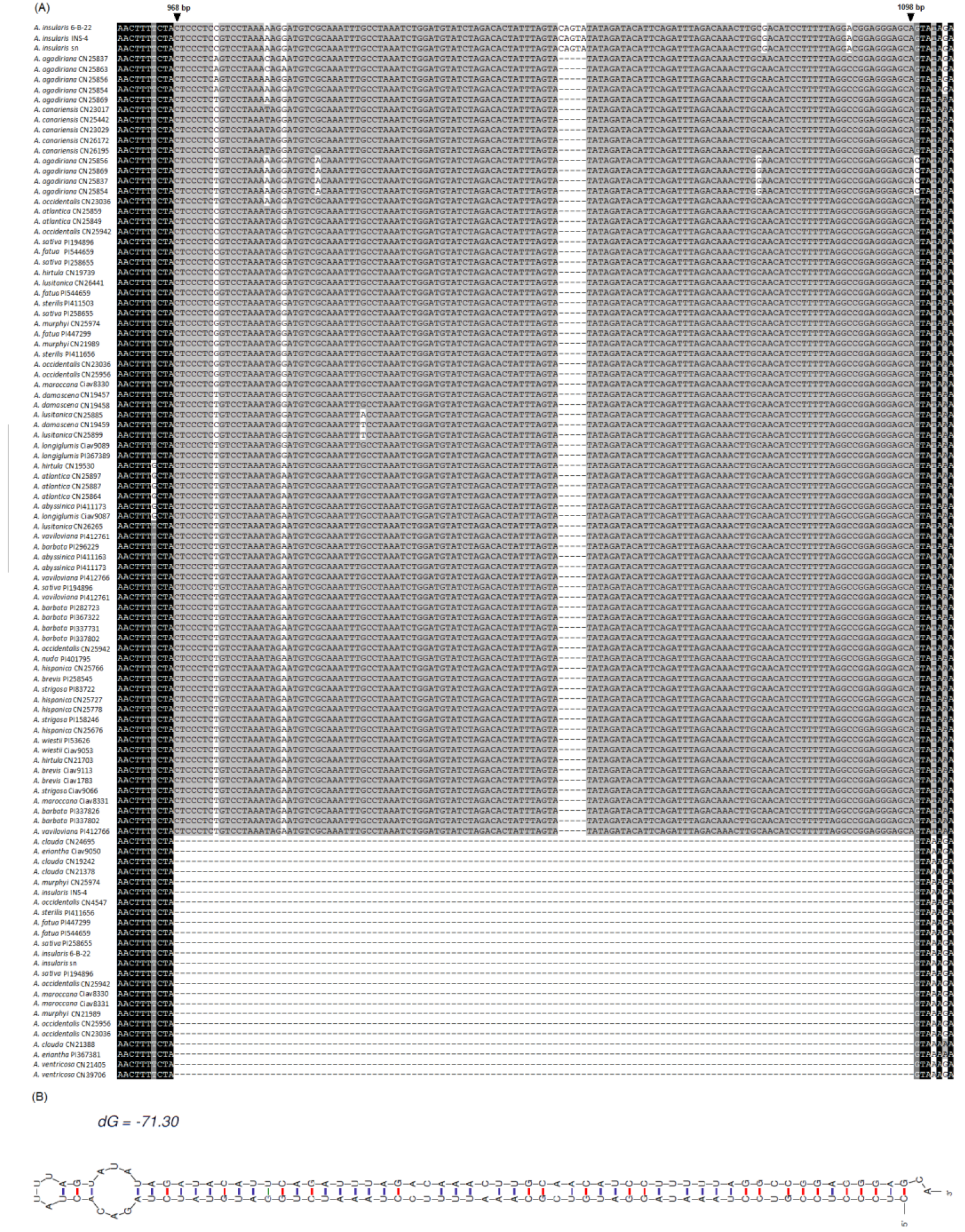
*pgk1* gene sequence analysis. (A) Partial alignment of the amplified *pgk1* gene of *Avena* species (B) Secondary structure of the deletion sequence between the A and C genomes.

### Phylogenetic analyses

The substitution plot for *pgk1* (Fig 2) indicated that the *pgk1* gene was not saturated and that it could be used for phylogenetic analysis. Phylogentic trees of 76 *Avena* accessions with the oat-like species *Trisetopsis turgidula* as outgroup were generated through maximum parsimony and Bayesian inference approaches on th non-redundant dataset. The parsimony analysis resulted in 80 equally parsimonioi trees (consistency index (CI) =0.632, retention index (RI) =0.954). BI analysis inferr an almost identical tree topology as the MP analysis, so the MP results were selected to describe this study (Fig 3).

**Fig 2.**
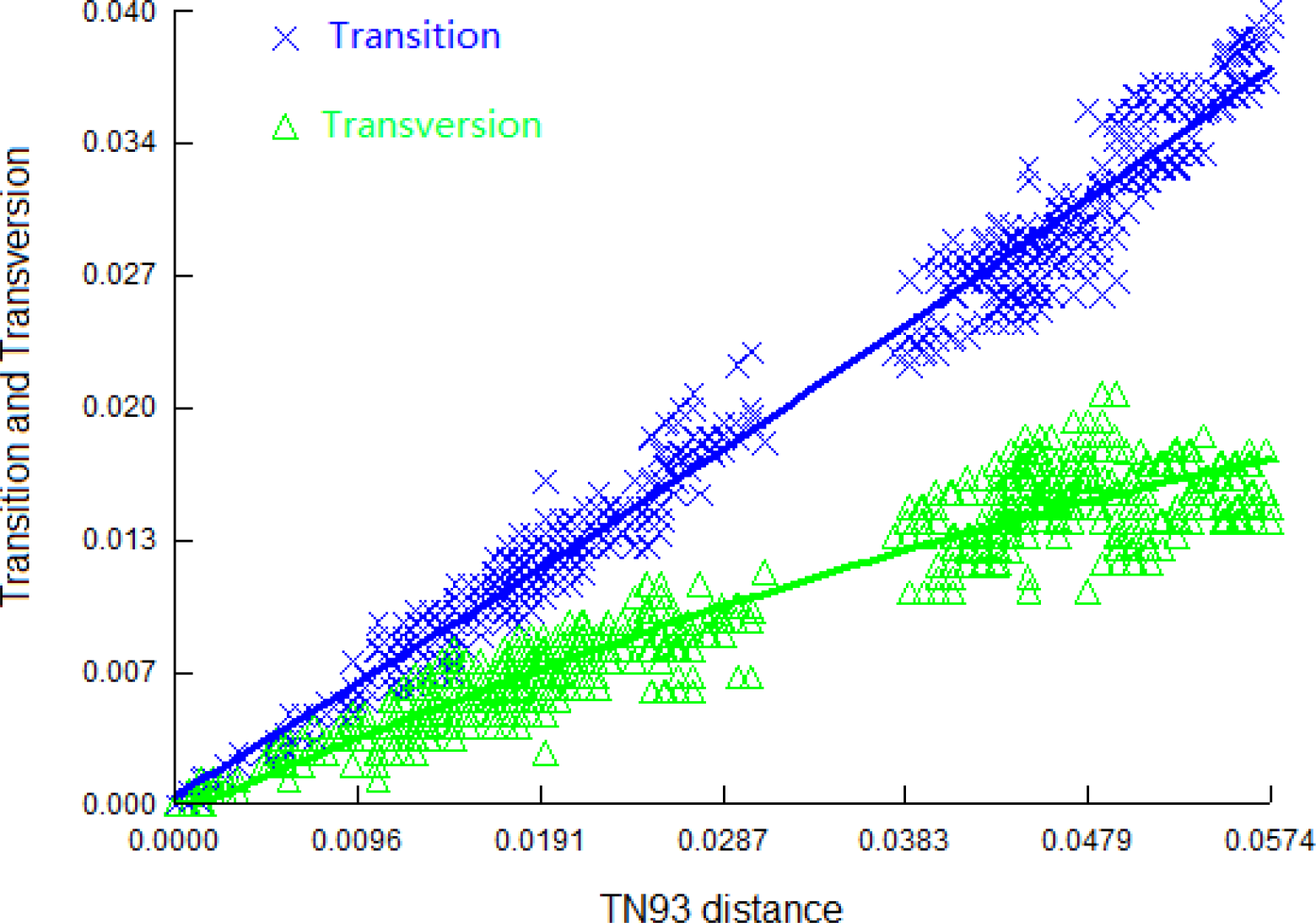
Saturation plot for transition and transversion of *pgk1* gene sequences. The crosses are the number of transition events; the triangles are the number of transversion events. The x axis shows the genetic distance based on the TN93 model; the y axis is the proportion of transitions or tansversions, which was calculated by using the number of transitions or transversions divided by the sequence length. The curves show the trends of the variance of transitions and transversions with the genetic distance increasing.

**Fig 3.**
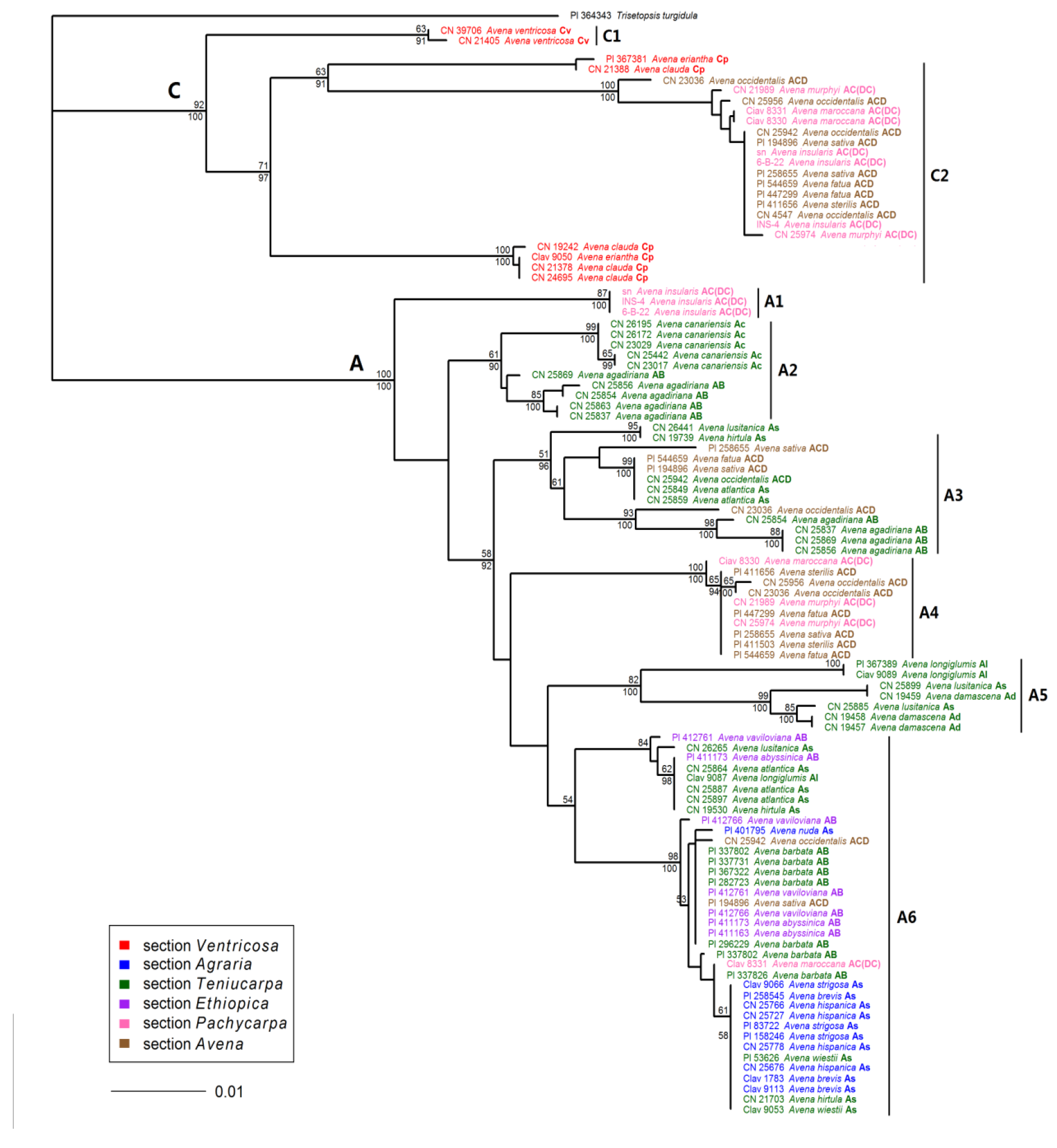
Maximum parsimony tree derived from *pgk1* sequence data. The tree was constructed using a heuristic search with TBR branch swapping. Numbers above and below the branches are bootstrap support (BS) values ≥50% and Bayesian posterior probability (PP) values ≥90%. Accession number, species name and haplome are indicated for each taxon.

Fig 3 shows that the *pgk1* gene sequences from 76 *Avena* accessions were split into two distinct clades with high BS (100% and 92%) and PP (100% and 100%) support. One clade contained all C-genome type sequences, hence referred to as the C genome clade. The other clade contained all sequences from the species carrying the A genome, henceforth, referred to as the A genome clade. The C genome clade was composed of two major subclades. All C_v_ genome diploids and two C_p_ genome diploid accessions formed the subclade C1 with 63% BS and 91% PP support, while subclade C2 included four C_p_ diploids accessions, seven AC(DC) genome tetraploid accessions and nine hexaploid accessions with 71% BS and 97% PP support. The *pgk1* gene sequences in the A genome clade were further split into six major subclades. The AC(DC) genome tetraploid species *A. insularis* was distinct from the other species, consequently forming a monophyletic clade (A1) with high BS (87%) and PP (100%) support. All five accessions of the A_c_ genome diploid species *A. canariensis* and one genome homoeologue of the AB genome tetraploid species *A. agadiriana* clustered together into subclade A2. Subclade A3 was composed of four accessions of the AB genome tetraploids *A. agadiriana*, five hexaploid accessions (A. *occidentalis* CN 23036 and CN 25942, *A. sativa* PI 194896 and PI 258655, *A. fatua* PI 544659) and four A_s_ genome diploid accessions ( *A. atlantica* CN25849 and CN 25859, *A. lusitanica* CN 26441, and *A. hirtula* CN 19739). One genome sequence of the AC(DC) genome tetraploids (without *A. insularis)* and the hexaploids formed a homogeneous clade (A4) that was separated from other species with high BS (100%) and PP (100%) support. The subclade A5 consisted of the A_d_ genome diploid *A. damascena,* the A_l_ genome diploid *A. longiglumis,* and the A_s_ genome diploid *A. lusitanica.* The remaining sequences from the A genome diploids and the AB genome tetraploids (without *A. agadiriana)* formed a relatively broader cluster A6, together with two hexaploid accessions (A. *sativa* PI 194896 and *A. occidentalis* CN 25942) and one AC (DC) genome tetraploid accession (A. *maroccana* CIav 8831).

Three groups of haplotypes of *pgk1* sequences were identified in five hexaploid accessions (A. *fatua* PI 544659, *A. occidentalis* CN 25942, CN 23036, and *A. sativa* PI 194896, PI 258655). These sequences fell into four subclades. One group clustered with the C genome diploids in subclade C2, and one group clustered with AC(DC) genome tetraploids in subclade A4. We hypothesize that these two types represent homoeologues from the C and D genomes, respectively. A third and fourth group fell into subclades A3 and A6. Since these two groups are highly separated, it is possible that they represent different A-genome events leading to different hexaploid lineages.

### Network analysis

To gain better insight into relationships within closely related lineages, MJ network reconstruction based on the haplotypes of *pgk1* sequences was employed. Due to the potential presence of recombination in the exon regions, the intron data was used for MJ network reconstruction. A total of 40 haplotypes were derived from 104 *pgk1* gene sequences (Fig 4). This low level of haplotype diversity demonstrates the high conservation of this gene within genus *Avena.* The MJ network recovered a nearly identical phylogenetic reconstruction to that based on the MP and BI trees, therefore we identified the clades from the MP results (Fig 3) within the MJ network (Fig 4). Based on the topology and frequency of haplotypes, the MJ network was split into two main groups. The two major groups representing two distinct types of haplotypes (A and C genomes) were distinguished due to the 131 bp insertion/deletion. Ten C genome haplotypes were observed, which were much less diverse than the 30 A genome haplotypes. The two main groups were further subdivided into clusters corresponding to the eight MP-based subclades discussed earlier. The only divergence was that the AC(DC) genome tetraploids *A. insularis,* which formed a separate clade (A1) in MP and BI trees, fell into together with the AB genome tetraploid *A. agadiriana* and the A_c_ genome diploid *A. canariensis* to form a relatively broad cluster in the MJ network (A1&A2).

**Fig 4.**
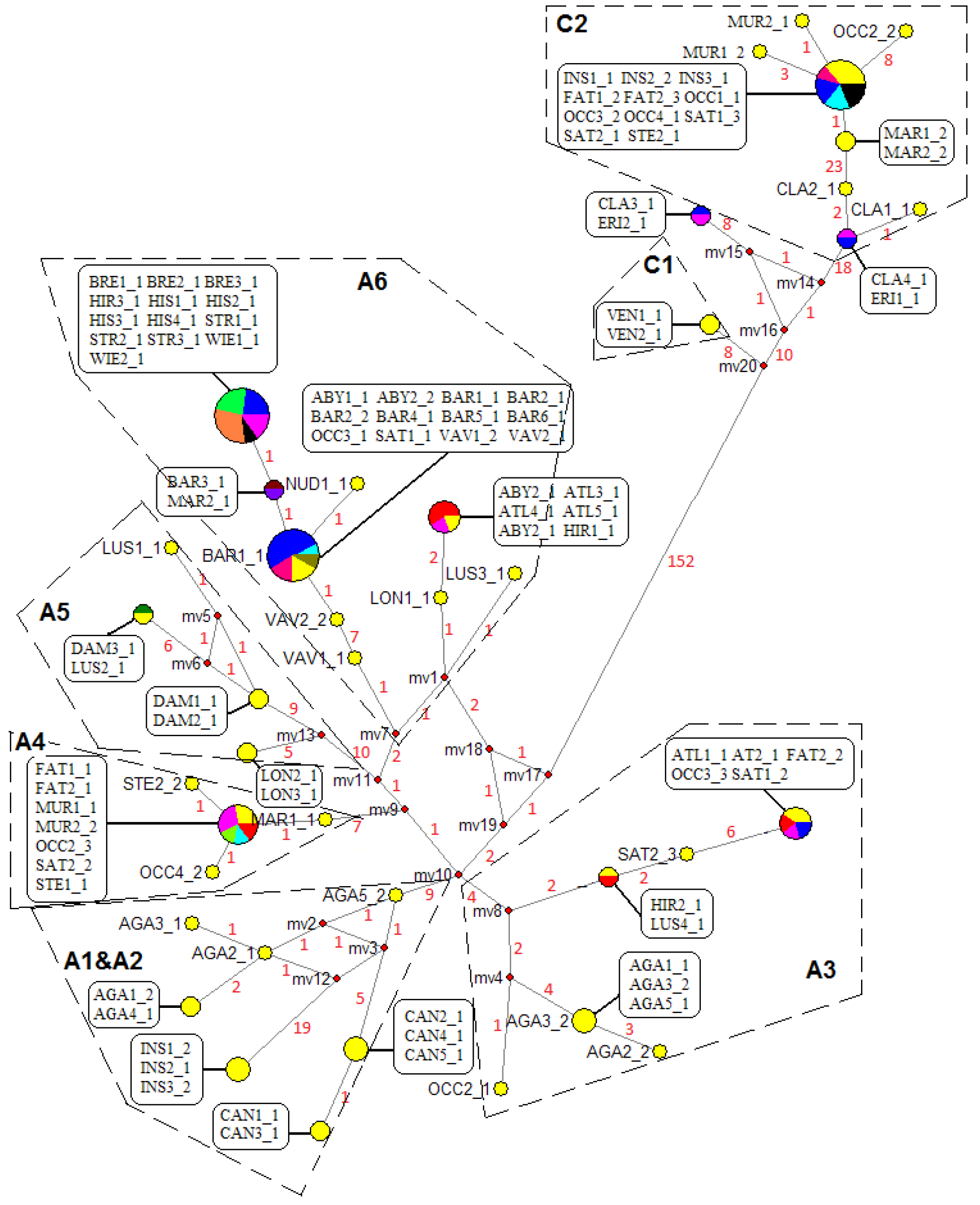
Median-joining networks based on 40 *pgk1* gene haplotypes of intron regions derived from 26 *Avena* species. Each circular node represents a single haplotype, with relative size being proportional to the frequency of that haplotype. Distinct colors in the same haplotype node represent different species sharing the same haplotype (colors are arbitrary). Median vectors (mv) represent the putative missing intermediates. Numbers along network branches indicate the number of bases involved in mutations between two nodes. Clusters (surrounded by dashed lines) are named based on clade names shown in the MP tree (Fig 3). Three-letter abbreviations of species names are listed in Table 1. The numbers immediately after each species abbreviation represent different accessions of the same species, and the number following the underscore identifies different haplotypes from the same accession.

## Discussion

### Two distinct diploid lineages exist in genus *Avena*

A significant 131 bp insert/deletion separated all *Avena* diploid species into two distinct groups representing the A and C genomes, respectively (Figs 1 and 4). These groups were also separated based on the MP or BI analysis that ignored gaps (Fig 3), indicating that the separation of A and C genomes is the most ancient major articulation in the genus *Avena,* a result that is consistent with most other literature [13, 14, 40]. MJ network analysis revealed that the C genome diploids have much lower levels of haplotype diversity than the A genome diploids. Within the C genome diploids, the C_p_ genome haplotypes were relatively more diverse than those of the C_v_ genome. These results might be explained by the geographic distribution of these species. The A genome diploids are distributed in a large region between latitude 20 and 40° N, while the C genome diploid species are restricted to a narrow territory along the Mediterranean shoreline [1]. The geographic distributions of the C genome diploid species are overlapping, but the range of the C_p_ genome diploid species is much broader than that of the C_v_ genome diploid species [41].

The A genome diploid species are the most diverse set of species in genus *Avena*, and chromosome rearrangements have occurred during the divergence of A-genomes from a common progenitor [41], resulting in the subdivision of the A genome into five types, of which we have investigated four. Our results showed that species with genome types A_c_, A_l_, and A_d_ formed groups that correspond well with previously reported structural differences. However, the A_s_ genome diploids appear to be much more diverse than previously reported, and are scattered into different subclades (Fig 3). Baum [1] divided all A_s_ genome diploids into two sections, section *Agraria* and section*Tenuicarpa.* All species of section *Agraria* have florets with a domesticated (non-shattering) base, whereas the other A_s_ species share relatively narrow spikelets. However, classification based on simple morphological traits is increasingly controversial. In this study, the A_s_ genome diploid species of section *Agraria* showed high degree of genetic homogeneity, consistently forming their own subclade A6, but other A_s_ genome species in section *Tenuicarpa* did not have their own subclade. *A. wiestii* showed a close relationship with the species of section *Agraria*, suggesting that it may be better-classified within that section. This result is in agreement with previous studies based on RAPD (Perchuk et al. 2002) and karyotypic comparisons (Badaeva et al. 2005). Accessions of the other two A_s_ genome species of section *Tenuicarpa (A. atlantica* and *A. hirtula)* were scattered into different subclades. These results were also observed in other studies (Peng et al. 2010, Yan et al. 2014). *A. lusitanica,* another A_s_ species of section *Tenuicarpa,* was diverged from other A_s_ species, but showed a close relationship to those with the A_d_ genome species *A. damascena.* This divergence has also been observed in many other studies [8, 9, 14, 40]. These, and other incongruences between morphological characters and genetic differences raise questions about appropriate taxonomical classifications among A_s_ genome species.

### The A_s_ and A_c_ genomes played roles in the AB tetraploid formation

Four recognized species have been proposed to have an AB genome composition. Of these, *A. barbata, A. abyssinica* and *A. vaviloviana* are grouped into a biological species known as the *barbata* group, while *A. agadiriana* is distinct [25, 42]. Our results confirmed the reported structural differences between these two groups (Fig 3). Two different *pgk1* gene sequences were detected from most of the AB genome tetraploids, supporting their allotetraploid origins. However, the genomes of *A.barbata* showed the least divergence, with only one of six *A. barbata* accessions providing multiple sequences, both of which were very similar. It seems that little divergence has occurred within the genome of *A. barbata* compared with that of *A. abyssinica* and *A.vaviloviana*, suggesting that *A. barbata* is the ancestral version of the species within the *barbata* group. This is supported by two lines of evidence. First, both *A. abyssinica* and *A.vaviloviana* are semi-domesticated forms that occur almost exclusively in Ethiopia, whereas the wild *A. barbata* are more geographically distributed, but can still be found close to the *abyssinica* and *vaviloviana* forms [43]. The second line of evidence was provided by FISH and Southern hybridization [17], which found some B chromosomes of *A. vaviloviana* are involved in inter-genomic translocations, while these rearrangements were not detected in *A. barbata.* There is little doubt that the A genome diploids have been involved in the formation of the *barbata* species. Some studies have suggested that both the A and B genomes of *barbata* species are diverged A_s_ genomes [16, 23, 44], while some others proposed that the B genome might have originated from other A genome diploid species [24, 25, 45]. In this study, both types of *pgk1* sequences detected from the *barbata* group showed high degree of genetic homogeneity with the A_s_ genome diploids (Fig 3), thus it was impossible to determine which type represents the A or B genome.

The recently discovered tetraploid species *A. agadiriana* was also proposed to have an AB genome composition because of its high affinity with *A. barbata* [23]. However, this designation has been questioned due to chromosomal divergences between *A. agadiriana* and the *barbata* species, as revealed by cytological studies [45, 46] and by molecular data [9, 13, 14]. In the current study, two distinct types of *pgk1* sequences were obtained in *A. agadiriana.* One copy clustered with the A_c_ genome species *A. canariensis*, whereas the other copy fell into cluster A3 with the A_s_ species *A. atlantica*, *A. hirtula*, *A. lusitanica*, and the hexaploids *A. occidentalis*, A. *fatua* and *A. sativa* (Fig 3). These results were in agreement with our previous studies based on nuclear *Acc1* gene [14] and GBS markers [9], and they support the proposal that *A. agadiriana* contains a different combination of A and/or B genomes from the *barbata* group, and that one of its two genomes originates from the A_c_ genome species *A. canariensis*, whereas the other one is closely related to the A_s_ species.

### The tetraploid species *A. maroccana* and *A. murphyi* are closely related to the hexaploids, while *A. insularis* is diverged

The other tetraploid group (*Avena* sect. *Pachycarpa*) contains three species, *A. maroccana, A. murphyi,* and the recently discovered *A. insularis.* Initial studies based on genomic in situ hybridization [47] supported an AC genome designation for these species. However, this designation has been challenged by FISH analysis, which has revealed that this set of tetraploid species, like the D chromosomes of the hexaploid oats, lacks a repetitive element that is diagnostic of the A genome [18]. This, together with other molecular evidence [14, 48] and our recent whole-genome analysis based on GBS markers [9], suggests that these tetraploid species contain the genome designated as D in hexaploid oats, and that they are more properly designated as DC genome species.

In the present study, two distinct *pgk1* homoeologues were detected in each of the three AC(DC) species, with each pair falling consistently into two clusters within the C and the A genome clades, respectively (Fig 3). The C-copy sequences of these tetraploids clustered consistently with the C-type homoeologues of the hexaploids, while the A/D genome homoeologues, with the exception of these from *A. insularis* and one sequence from *A. maroccana* (CIav 8331) fell into subclade A4 along with a set of sequences from the hexaploid oats (Fig 3). Considering that the other *pgk1* gene sequences from the hexaploid oats clustered with the C or A genome diploids, we deduced that the sequences falling in subclade A4 must represent the D genome homoeologues of the hexaploids and of the AC(DC) species *A. maroccana* and *A. murphyi.* This result is not fully consistent with our previous GBS study: although *A. maroccana* and *A. murphyi* were very similar to hexaploid oat and were designated as DC genomes, our GBS work suggested that *A. insularis* was also a DC genome that was even more similar to the hexaploids [9]. Examining the existing literature, all three of these tetraploid species have variously been considered as the tetraploid ancestor of the hexaploids [4, 9, 49]. In view of the genome structure of these tetraploids [24, 50] and the meiotic chromosome paring of their interspecific hybrids [51], all of these tetraploids are proposed to have diverged from a common ancestral tetraploid after the occurrence of some large chromosome rearrangements [24, 50]. However, it cannot be ruled out that these tetraploids might have originated independently from different diploid ancestors, since they have shown close relationships with different diploid species [40]. Interestingly, in network analysis (Fig 4), the A/D-type homoeologues of *A. insularis* fell into a group with the A_c_ genome species *A. canariensis* and the AB genome species *A. agadiriana.* In fact, previous studies have revealed that *A. canariensis* is closely related to the DC genome tetraploids [15]. These results suggest a possibility that *A. canariensis* was involved in contributing an early version of a D genome in all three AC(DC) genome tetraploids. Nevertheless, we do not have an explanation for why the D genome copy of *pgk1* in *A. insularis* could have diverged so far from the version found in the hexaploids, especially since the C genome copies remain identical.

### The genome origins of the hexaploid species

It is now generally accepted that two distinct steps were involved in the evolution of hexaploid oats. The first step would have been the formation of a DC genome hybrid from ancestral D and C genome diploids, followed by doubling of the chromosomes to form an allotetraploid. The second step would have involved hybridization of a DC tetraploid with a more recent A genome diploid, followed by doubling of the triploid hybrid [9, 13].

The diploid progenitor of the hexaploid C genome was probably restricted to the narrow geographic range where the three extant C genome diploids are distributed. However, numerous inter-genomic translocations among hexaploid chromosomes [9, 11, 52, 53]have deceased the homology between the C genome diploids and the hexaploid C genome, making the identification of the C genome donor of the hexaploids challenging. In this study, the C_p_ genome species shared the highest degree of genetic similarity with both the DC genome tetraploids, as well as with the hexaploids, leading us to conclude that a C_p_ genome species was the C genome donor of the polyploids. This conclusion is supported by other evidence from nuclear genes [13, 54]. This is important, since it was recently demonstrated that the maternal tetraploid and hexaploid genomes originated from an A genome species, not from a C genome species [55], rendering comparisons to the C_v_ vs C_p_ maternal genomes irrelevant in determining the origin of the nuclear C genome in the hexaploids.

The A genome origin of the hexaploids remains a matter of debate, and many A genome diploids have been suggested as putative diploid progenitors, as summarized by Peng et al [13]. FISH analysis showed that an A_s_-specific DNA repeat was restricted to the A_s_ and A_l_ genomes, as well as the hexaploid A genome [18]. In this study, a close relationship between the A_s_ genome diploid *A. atlantica* was observed for some hexaploid haplotypes in the phylogenetic tree (Fig 3) and the MJ network (Fig 4). An *A. atlantica* genome origin is consistent with previous studies based on IGS-RFLP analysis [12] and the *ppcB1* gene [40]. However, there is evidence in our work that some hexaploids may have an alternate A genome origin, closer to the *Agraria* section of A_s_ diploids. The presence of multiple A genome origins could explain variable results that have been reported in studies of hexaploid phylogeny.

In this study, strong evidence is presented for a D genome origin in the tetraploids *A. maroccana* and *A. murphyi* (Figs 3-4). However, these D genome sequences did not show a close relationship with any diploid species investigated in this study. Other than the discrepancy with *A. insularis,* this result is consistent with our recent GBS study [9]. One factor that may hinder the discovery of a D genome progenitor is the presence of inter-genomic translations among all three genomes in the hexaploid [9, 53]. Two hexaploid accessions *(A.occidentalis* CN 25942 and *A. sativa* PI 194896) did not contribute haplotypes that clustered with the putative D genome sequences (Subclade A4 in Fig 3). Although this may be a result of incomplete sampling, it may also result from inter-genomic translations that have duplicated or eliminated copies of *pgk1*.

In conclusion, this is the most comprehensive study to date that investigates a phylogeny in genus *Avena* using a single informative nuclear gene. It confirms or clarifies most previous work, and presents strong evidence in support of a working hypothesis for the origin of hexaploid oat. However, many questions still remain, and these will be best addressed through further studies involving multiple nuclear genes or whole genomes. We are collaborating on work that will provide exome-based gene diversity studies, but this work will require complete hexaploid reference sequences before it can be properly analyzed. Such reference sequences are currently in progress, so the next few years may see a revolution in our understanding of *Avena* phylogeny. Nevertheless, as many researcher in this field are aware, the polyploid species in this genus have experienced extensive chromosome rearrangement, which will continue to complicate phylogenetic studies. It may even be necessary to generate a pan-genome hexaploid reference sequence before definitive statements can be made.

## Acknowledgements

We are very grateful to the Plant Gene Resources of Canada (PGRC), the National Small Grains Collection, Agriculture Research Service, United States Department of Agriculture (USDA, ARS) and Dr. Rick Jellen, Brigham Young University providing seed materials. We also thank the anonymous reviewers for the useful comments on this manuscript.

## Reference

1. Baum BR. Oats: wild and cultivated. A monograph of the genus Avena L.(Poaceae): Minister of Supply and Services; 1977.

2. Baum BR, Fedak G. Avena atlantica, a new diploid species of the oat genus from Morocco. Canadian Journal of Botany. 1985;63(6): 1057–1060.

3. Baum BR, Fedak G. A new tetraploid species of *Avena* discovered in Morocco. Canadian Journal of Botany. 1985;63(8): 1379–1385.

4. Ladizinsky G. A new species of *Avena* from Sicily, possibly the tetraploid progenitor of hexaploid oats. Genetic Resources and Crop Evolution. 1998;45(3): 263–269.

5. Lin L, Liu Q. Geographical distribution of *Avena* L.(Poaceae). Journal of Tropical & Subtropical Botany. 2015;2: 111–122.

6. Rajhathy T, Thomas H. Cytogenetics of oats (Avena L.): Genetics Society of Canada; 1974.

7. Thomas H. Cytogenetics of Avena. In: Marshall HG, Sorrells ME, editors. Oat Science and Technology. Agronomy Monograph. Madison, WI: American Society of Agronomy, Crop Science Society of America; 1992. pp. 473–507.

8. Yan H, Martin SL, Bekele WA, Latta RG, Diederichsen A, Peng Y, et al. Genome size variation in the genus *Avena*. Genome. 2016;59(3): 209–220.

9. Yan H, Bekele WA, Wight CP, Peng Y, Langdon T, Latta RG, et al. High-density marker profiling confirms ancestral genomes of *Avena* species and identifies D-genome chromosomes of hexaploid oat. Theoretical and Applied Genetics. 2016;129(11): 2133–2149.

10. Rajhathy T, Thomas H. Chromosomal differentiation and speciation in diploid *Avena*. III. Mediterranean wild populations. Canadian Journal of Genetics and Cytology. 1967;9(1): 52–68.

11. Chen Q, Armstrong K. Genomic in situ hybridization in *Avena sativa*. Genome. 1994;37(4): 607–612.

12. Nikoloudakis N, Katsiotis A. The origin of the C-genome and cytoplasm of *Avena* polyploids. Theoretical and Applied Genetics. 2008;117(2): 273–281.

13. Peng Y-Y, Wei Y-M, Baum BR, Yan Z-H, Lan X-J, Dai S-F, et al. Phylogenetic inferences in *Avena* based on analysis of *FL* intron2 sequences. Theoretical and Applied Genetics. 2010;121(5): 985–1000.

14. Yan H-H, Baum BR, Zhou P-P, Zhao J, Wei Y-M, Ren C-Z, et al. Phylogenetic analysis of the genus *Avena* based on chloroplast intergenic spacer psbA-trnH and single-copy nuclear gene Acc1. Genome. 2014;57(5): 267–277.

15. Fominaya A, Vega C, Ferrer E. Giemsa C-banded karyotypes of *Avena* species. Genome. 1988;30(5): 627–632.

16. Katsiotis A, Hagidimitriou M, Heslop-Harrison JS. The close relationship between the A and B genomes in *Avena* L. (Poaceae) determined by molecular cytogenetic analysis of total genomic, tandemly and dispersed repetitive DNA sequences. Annals of Botany. 1997;79(2): 103–109.

17. Irigoyen M, Loarce Y, Linares C, Ferrer E, Leggett M, Fominaya A. Discrimination of the closely related A and B genomes in AABB tetraploid species of *Avena*. Theoretical and Applied Genetics. 2001;103(8): 1160–1166.

18. Linares C, Ferrer E, Fominaya A. Discrimination of the closely related A and D genomes of the hexaploid oat *Avena sativa* L. Proceedings of the National Academy of Sciences. 1998;95(21): 12450–12455.

19. Chew P, Meade K, Hayes A, Harjes C, Bao Y, Beattie AD, et al. A study on the genetic relationships of *Avena* taxa and the origins of hexaploid oat. Theoretical and Applied Genetics. 2016;129(7): 1405–1415.

20. Lináres C, Gonzalez J, Ferrer E, Fominaya A. The use of double fluorescence in situ hybridization to physically map the positions of 5S rDNA genes in relation to the chromosomal location of 18S-5.8S-26S rDNA and a C genome specific DNA sequence in the genus Avena. Genome. 1996;39(3): 535–542.

21. Leggett J, Markhand G, editors. The genomic structure of *Avena* revealed by GISH. Proceedings of the Kew Chromosome Conference IV; 1995.

22. Baum BR, Rajhathy T, Sampson DR. An important new diploid *Avena* species discovered on the Canary Islands. Canadian Journal of Botany. 1973;51(51): 759–762.

23. Leggett JM, Thomas H. Oat evolution and cytogenetics. In: Welch RW, editor. The Oat Crop World Crop Series: Springer, Dordrecht; 1995. pp. 120–149.

24. Fominaya A, Vega C, Ferrer E. C-banding and nucleolar activity of tetraploid *Avena* species. Genome. 1988;30(5): 633–638.

25. Badaeva E, Shelukhina OY, Goryunova S, Loskutov I, Pukhalskiy V. Phylogenetic relationships of tetraploid AB-genome *Avena* species evaluated by means of cytogenetic (C-banding and FISH) and RAPD analyses. Journal of Botany. 2010; 2010.

26. Fan X, Sha LN, Zeng J, Kang HY, Zhang HQ, Wang XL, et al. Evolutionary dynamics of the *pgk1* gene in the polyploid genus *Kengyilia* (Triticeae: Poaceae) and its diploid relatives. Plos One. 2012;7(2): e31122.

27. Sha LN, Fan X, Wang XL, Dong ZZ, Zeng J, Zhang HQ, et al. Genome origin, historical hybridization and genetic differentiation in *Anthosachne australasica* (Triticeae; Poaceae), inferred from chloroplast *rbc*L, *trn*H-*psb*A and nuclear Acc1 gene sequences. Ann Bot. 2017;119(1): 95–107.

28. Huang S, Sirikhachornkit A, Faris JD, Su X, Gill BS, Haselkorn R, et al. Phylogenetic analysis of the acetyl-CoA carboxylase and 3-phosphoglycerate kinase loci in wheat and other grasses. Plant Molecular Biology. 2002;48(5-6): 805–820.

29. Chen Q, Kang HY, Fan X, Wang Y, Sha LN, Zhang HQ, et al. Evolutionary history of *Triticum petropavlovskyi* Udacz. et Migusch. inferred from the sequences of the 3-Phosphoglycerate kinase gene. Plos One. 2013;8(8): e71139.

30. Ladizinsky G. *Avena prostrata*: a new diploid species of oat. Israel J Bot. 1971: 297–301.

31. Doyle J. A rapid DNA isolation procedure for small quantities of fresh leaf tissue. Phytochemical Bulletin. 1987;19(1): 11–15.

32. Thompson JD, Higgins DG, Gibson TJ. CLUSTAL W: improving the sensitivity of progressive multiple sequence alignment through sequence weighting, position-specific gap penalties and weight matrix choice. Nucleic Acids Research. 1994;22(22): 4673–4680.

33. Xia X. DAMBE5: A comprehensive software package for data analysis in molecular biology and evolution. Molecular Biology and Evolution. 2013;30(7): 1720–1728.

34. Swofford DL. PAUP: Phylogenetic analysis using parsimony (and other metods). Version 4.0b.10. Sunderland: Sinauer Associates. 2003.

35. Felsenstein J. Confidence limits on phylogenies: an approach using the bootstrap. Evolution. 1985;39(4): 783–791.

36. Huelsenbeck JP, Ronquist F. MRBAYES: Bayesian inference of phylogenetic trees. Bioinformatics. 2001;17(8): 754–755.

37. Bandelt HJ, Forster P, Röhl A. Median-joining networks for inferring intraspecific phylogenies. Molecular Biology and Evolution. 1999;16(1): 37–48.

38. Kilian B, Özkan H, Deusch O, Effgen S, Brandolini A, Kohl J, et al. Independent wheat B and G genome origins in outcrossing *Aegilops* progenitor haplotypes. Molecular Biology and Evolution. 2007;24(1): 217–227.

39. Pond SLK, Frost SDW, Muse SV. HyPhy: hypothesis testing using phylogenies. Bioinformatics. 2005;21(5): 676–679.

40. Liu Q, Lin L, Zhou X, Peterson PM, Wen J. Unraveling the evolutionary dynamics of ancient and recent polyploidization events in *Avena* (Poaceae). Scientific Reports. 2017;7: 41944.

41. Loskutov IG, Rines HW. Avena. In: Kole C, editor. Wild crop relatives: genomic and breeding resources: Springer; 2011. pp. 109–183.

42. Drossou A, Katsiotis A, Leggett JM, Loukas M, Tsakas S. Genome and species relationships in genus *Avena* based on RAPD and AFLP molecular markers. Theoretical and Applied Genetics. 2004;109(1): 48–54.

43. Ladizinsky G. Studies in Oat Evolution: Springer Berlin Heidelberg; 2012.

44. Holden JHW. Species relationships in the Avenue. Chromosoma. 1966;20: 75–124.

45. Shelukhina OY, Badaeva ED, Brezhneva TA, Loskutov IG, Pukhalsky VA. Comparative analysis of diploid species of *Avena* L. using cytogenetic and biochemical markers: *Avena canariensis* Baum et Fedak and *A. longiglumis* Dur. Russian Journal of Genetics. 2008;44(6): 694–701.

46. Jellen EN, Gill BS. C-banding variation in the Moroccan oat species *Avena agadiriana* (2n=4×=28). Theoretical and Applied Genetics. 1996;92(6): 726–732.

47. Jellen EN, Gill BS, Cox TS. Genomic in situ hybridization differentiates between A/D- and C-genome chromatin and detects intergenomic translocations in polyploid oat species (genus *Avena*). Genome. 1994;37(4): 613–618.

48. Oliver RE, Jellen EN, Ladizinsky G, Korol AB, Kilian A, Beard JL, et al. New Diversity Arrays Technology (DArT) markers for tetraploid oat (*Avena magna Murphy* et Terrell) provide the first complete oat linkage map and markers linked to domestication genes from hexaploid *A. sativa* L. Theoretical & Applied Genetics. 2011;123(7): 1159–1171.

49. Ladizinsky G, Zohary D. Notes on species delimination, species relationships and polyploidy in *Avena* L. Euphytica. 1971;20(3): 380–395.

50. Shelukhina OY, Badaeva ED, Loskutov IG, Pukhal’sky VA. A comparative cytogenetic study of the tetraploid oat species with the A and C genomes: *Avena insularis, A. magna*, and *A. murphyi*. Russian Journal of Genetics. 2007;43(6): 613–626.

51. Ladizinsky G. Cytogenetic relationships between *Avena insularis* (2n=28) and both *A. strigosa* (2n=14) and *A. murphyi* (2n=28). Genetic Resources and Crop Evolution. 1999;46(5): 501–504.

52. Irigoyen ML, Linares C, Ferrer E, Fominaya A. Fluorescence in situ hybridization mapping of *Avena* sativa L. cv. SunII and its monosomic lines using cloned repetitive DNA sequences. Genome. 2002;45(6): 1230–1237.

53. Sanz MJ. A new chromosome nomenclature system for oat (*Avena sativa L.* and *A. byzantina* C. Koch) based on FISH analysis of monosomic lines. Theoretical and Applied Genetics. 2010;121(8): 1541–1552.

54. Cheng DW, Armstrong KC, Drouin G, Mcelroy A, Fedak G, Molnar SD. Isolation and identification of *Triticeae* chromosome 1 receptor-like kinase genes (*Lrk10*) from diploid, tetraploid, and hexaploid species of the genus *Avena*. Genome. 2003;46(1): 119–127.

55. Fu YB. Oat evolution revealed in the maternal lineages of 25 *Avena* species. Scientific Reports. 2018;8(1): 4252.

